# Time dependence of cellular responses to dynamic and complex strain fields

**DOI:** 10.1101/286625

**Authors:** Sophie Chagnon-Lessard, Michel Godin, Andrew E. Pelling

## Abstract

Exposing cells to an unconventional sequence of physical cues can reveal subtleties of cellular sensing and response mechanisms. We investigated the mechanoresponse of cyclically-stretched fibroblasts under a spatially non-uniform strain field which was subjected to repeated changes in stretching directions over 55 hours. A polydimethylsiloxane microfluidic stretcher array optimized for complex staining procedures and imaging was developed to generate biologically relevant strain and strain _gradient_ amplitudes. We demonstrated that cells can successfully reorient themselves repeatedly, as the main cyclical stretching direction is consecutively switched between two perpendicular directions every 11 hours. Importantly, from one reorientation to the next, the extent to which cells reorient themselves perpendicularly to the local strain direction progressively decreases, while their tendency to align perpendicularly to the strain gradient direction tends to increase. We demonstrate that these results are consistent with our finding that cellular responses to strains and strain gradients occur on two distinct time scales, the latter being slower. Overall, our results reveal the absence of major irreversible cellular changes that compromise the ability to sense and reorient to changing strain directions under the conditions of this experiment. On the other hand, we show how the history of strain field dynamics can influence the cellular realignment behavior, due to the interplay of complex time-dependent responses.

## INTRODUCTION

The use of lab-on-a-chip^1,2^ strategies allows mimicking complex in vivo physical cues and thus can facilitate the study of cellular behaviors in specific microenvironments. Cells in living organisms are constantly subjected to various mechanical forces such as compression and stretching (mechanical strains) due, for example, to muscle contractions, fluid pressure, pumping of the heart, and breathing. Such mechanical signals are known to be involved in the regulation of many fundamental biological processes such as cell migration, division, differentiation, and apoptosis^3–6^. Another example extensively studied to further our understanding of mechanotransduction processes is the cellular reorientation induced by cyclic stretching. More specifically, cyclically stretched cells were shown to exhibit elongation^7,8^, alignment perpendicular to the stretching direction^9–15^, and more recently fibroblasts were also demonstrated to exhibit strain gradient avoidance^16^.

In vivo microenvironments are known to impose an array of time-dependent mechanical strains on cellular systems. For example, some studies have highlighted the role of complex mechanoresponse pathways related to embryonic development, or in articular cartilage development and degeneration. In these cases, not only are the mechanical forces known to play a key role^17–19^, but they can be non-uniform and rapidly changing. There are also contexts in which the stiffness is perturbed locally, thus inducing strain gradients, for instance following the insertion of coronary artery stents^20^. In general, the complex microenvironment in which cells evolve in vivo gives rise to non-uniform, time-varying, anisotropic strain fields^21–24^. Importantly, in our previous work^16^, we have clearly demonstrated that cells have an ability to sense and respond to both strain and strain gradients. Yet, the large majority of previous cyclic stretching studies have focused on stretch patterns with nearly-uniform strain amplitude and direction^8–11,13,15^. Although the present study does not aim to directly mimic a specific in vivo condition, it investigates the effect of biologically relevant strain fields with artificial discontinuous events, considering that sudden mechanical alterations can occur in living organisms. By studying cell responses to an unconventional sequence of mechanical cues we aimed to reveal subtle aspects of their mechanosensitivity. In particular, it is poorly understood if cells would continue to exhibit the expected reorientation response if the strain field was repeatedly altered multiple times over the course of several days. Moreover, key insights on subtle cellular mechanisms can be gained by determining whether the cellular response to strain remains constant or if it changes over time with an increasing number of reorientation events.

This work investigates the influence of alternating the stretching axis of a complex strain field on the degree of cell alignment with the strain direction, the degree of cell alignment with the strain gradient direction, and the cell morphology. Sudden changes in stretching directions are achieved by employing a biaxial device which allows independent control over perpendicular stretching directions. In this context, we developed a highly versatile polymethylsiloxane (PDMS) microfluidics device integrating an array of stretchable membranes. In particular, it can be made open-top at any point during a given experiment to facilitate imaging, circumventing disadvantages of typical microfluidic stretchers. Specifically, it enables complex staining procedures with minimal cellular perturbations, as well as near-diffraction limited imaging.

Here, we use human foreskin fibroblasts (HFF) as a model system as we have previously shown that they are acutely sensitive to both strain and strain gradients^16^, thus being ideal for exploratory work. In the present study, we demonstrate that HFF cells possess the ability to reorient themselves when the imposed strain direction is changed, retaining this ability for at least five dramatic changes in strain direction. Our comprehensive analysis shows that the orientation distribution does vary with changes in stretching direction. Namely, we observed that the cells become progressively less oriented with the strain direction and conversely their alignment with the strain gradient direction improves as the experiment progresses. Furthermore, we show that this behavior is driven by a fascinating ability of these cells to respond to two separate, but simultaneous, mechanical cues over different time scales. Such complexity exists in vivo but is not widely reflected in fundamental studies as of yet due to the relatively few number of biaxial devices which allow for the production of complex strain fields. This study clearly highlights the importance of studying cellular behaviors in increasingly complex on-chip microenvironments.

## RESULTS AND DISCUSSION

### Microfluidic stretcher design

Microfluidic devices offer many advantages over manual methods by enabling automated on-chip sample handling capabilities, decreasing contamination risks, reducing reagent quantities, and enabling high fabrication reproducibility^25^. In principle, they also have the potential to enable high-throughput screening experiments. An often-overlooked advantage of microstretchers over macrostretchers in certain studies is that they naturally produce non-uniform strain fields with biologically relevant strain gradients, while maintaining the strain magnitude to desirably low levels^26^ (see Supporting Information). This enables the study of mechanical cues which better mimics the complexity of the cell microenvironment in vivo. However, in applications where cells need to be cultured within the chip, such as for cyclic-stretching experiments, numerous experimental challenges^27,28^ (nonuniform cell seeding, clogging, apparition of air bubbles, accumulation of undesired residues, etc) often impact the reliability of the experimental approach. Microfluidic devices also have disadvantages in the context of cell immunofluorescence microscopy. The first is the inherent difficulty in performing complex and sensitive straining procedures without compromising the integrity of the cell components (due to fluid shear stress^29^). The second issue is the reduction in imaging resolution which results from the optical aberrations introduced by the presence of PDMS in the optical path^30^.

To address these challenges, we built upon our previous design^31^ as well as those of others^32,33^ and we developed the 2×2 microstretcher array which working principle is summarized in Fig. 1. This new design allowed the study of ∼4×200 cells per array, leading to robust measurements enabling the detection of subtle changes which could have remained unexposed without such parallelization. Cyclic stretching is achieved by using vacuum chambers that repeatedly deform the walls surrounding a suspended membrane on which cells are adhered. The throughput and ease-of-use were considerably increased by scaling it into an array and most importantly by allowing the microdevice to be easily made open-top at any point over the course of an experiment, i.e. when the microfluidic properties are no more required or beneficial. In several instances, it will be desirable to preserve the microfluidic nature of the device over the course of the actual experiment but to convert it into an open-top device before staining and imaging.

**Figure 1.**
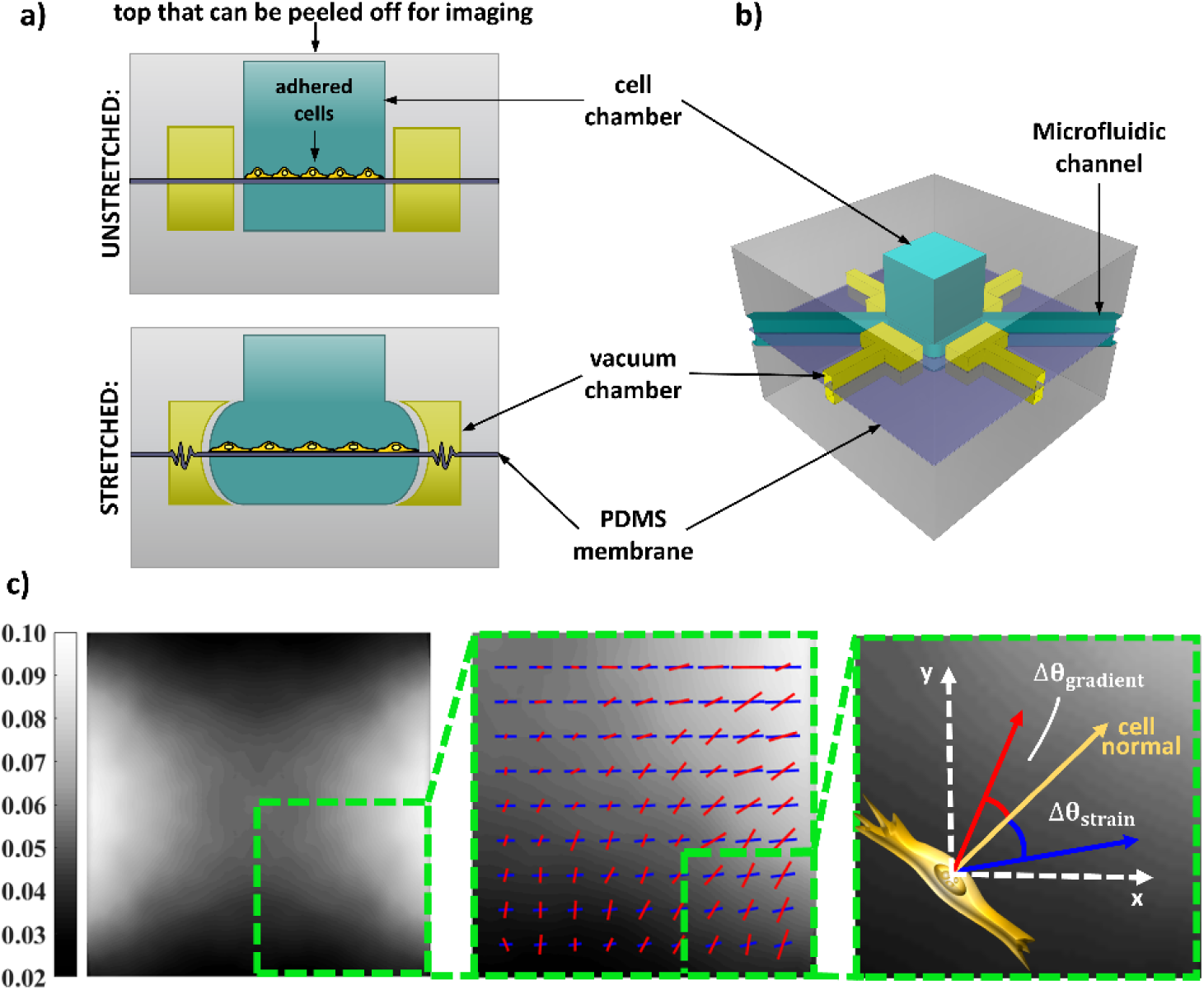
Microfluidics stretcher working principle, membrane strain field map and angle convention. **a)** Side view of the design presented to show the unstretched (top) and the stretched (bottom) states. The schematic is not to scale and has been adapted from previous work^26^. **b)** Three-dimensional view, partially transparent to show the design. **c)** Left: Colormap of the experimentally determined maximum principal strain amplitude ε1(x,y) for x-stretching. The full membrane is shown (1.6 mm × 1.6 mm). Middle: The bottom right corner of the membrane is zoomed such that 25% of the total area is displayed. A representation of the amplitude (line length) and direction (line orientation) of the maximum principal strain (blue lines) and its gradient (red lines) is also presented. Right: The bottom right corner is further zoomed in to depict a fictive cell (yellow) and the angle convention used throughout the article. The red arrow represents the gradient direction and the blue is the strain direction. The angle difference between the cell normal and the gradient direction is identified as Δθ_gradient_ while the angle difference between the cell normal and the strain direction is given by Δθstrain.

Previous pneumatic-based microfluidic stretchers were generally permanently closed^31–34^. The general approach used here consists instead in extending the cell chamber height in such a way as to leave only a thin PDMS layer to act as the top sealant. When properly fabricated, this thin layer (on the order of 200 μm thick) can be easily peeled off under a microscope with tweezers. It is worth noting that the ability to switch from a closed-top to an open-top microfluidic device is of general interest and has been addressed previously in other contexts by using a (non-PDMS) lid^35,36^. The all-PDMS strategy that we present here is an attractive alternative due to its fabrication and utilization simplicity, as well as its robustness with respect to eliminating leakage risks. Although disadvantageous for imaging, conserving the microfluidic properties can be highly beneficial in various studies investigating the combined effect of different stimuli, for instance flow shear stress or time-varying chemical concentrations, together with the underlying substrate strain^37–39^. The main fabrication steps are presented in the Materials and Methods section.

### Strain field characterization and device imaging performance

The membrane’s strain field generated in the microdevice is presented in Fig. 1c (for X-stretching). It is based on the analysis of the fluorescent beads displacements as explained in the Materials and Methods section. Importantly, our device’s geometry and scale have the advantage of naturally producing smoothly-varying strain gradients (from 0 to 14% mm^−1^) of significant and biologically relevant magnitudes^24^ while maintaining the strain amplitude relatively low (2 to 10%), i.e. also biologically relevant to optimally induce the cellular responses ^9,10,12^. Moreover, our design produces a strain field for which the position dependent principal strain direction is largely decoupled from that of the gradient. This is an essential condition to properly study their respective effects.

Efforts have been put recently in developing microfabrication strategies to allow high resolution microscopy in microfluidic devices by minimizing the optical path through PDMS which is known to introduce aberrations preventing diffraction-limited imaging^30,40^. Our approach presents an effective solution for microstretchers. Standard all-PDMS devices usually comprise >1mm thick layers to ensure structural integrity and proper external tubing connections^31^. Fig. 2 demonstrates the imaging capabilities of our hybrid microdevice once it has been made open-top. The image profile of a fluorescent bead obtained within the open-top chamber is compared with that obtained when a similar bead is covered with a ∼800 μm thick PDMS layer inserted in the light path (Fig. 2a). In average, the lateral full width at half maximum (FWHM) resolution degrades by (30 ± 3)% with the insertion of the PDMS layer (Fig. 2b). This loss of lateral resolution is in qualitative agreement with that measured by Tonin *et al*., who used an oil-immersion objective with which they actually observed an even more detrimental effect despite testing thinner layers^30^. Importantly, the spread in the distribution of lateral FWHMs shown in Fig. 2b is also significantly larger in the closed-top case, rendering deconvolution-based image analysis strategies less reliable^41^. Axially, we measured an average four-fold increase of the bead’s profile width (Fig. 2c), demonstrating a dramatic decrease in one’s ability to resolve features depth-wise in closed-top PDMS devices. Finally, we note that open-top imaging is also beneficial with respect to photobleaching since lower laser excitation intensities are required (∼five-fold in our case) to achieve similar peak signal intensities. The overall deterioration in resolution (in particular axially) can be attributed primarily to the introduction of spherical aberrations which result from refractive index mismatch^42–44^. Additional explanation is provided in the Supplementary Information (SI). Overall, our results show the advantage of using our design over a traditional closed-top microdevice. To illustrate the relevance of this high imaging capability in the context of cell biology, Fig. 2d shows the actin filaments and the focal adhesions of an HFF cell imaged in our device via immunofluorescence staining.

**Figure 2.**
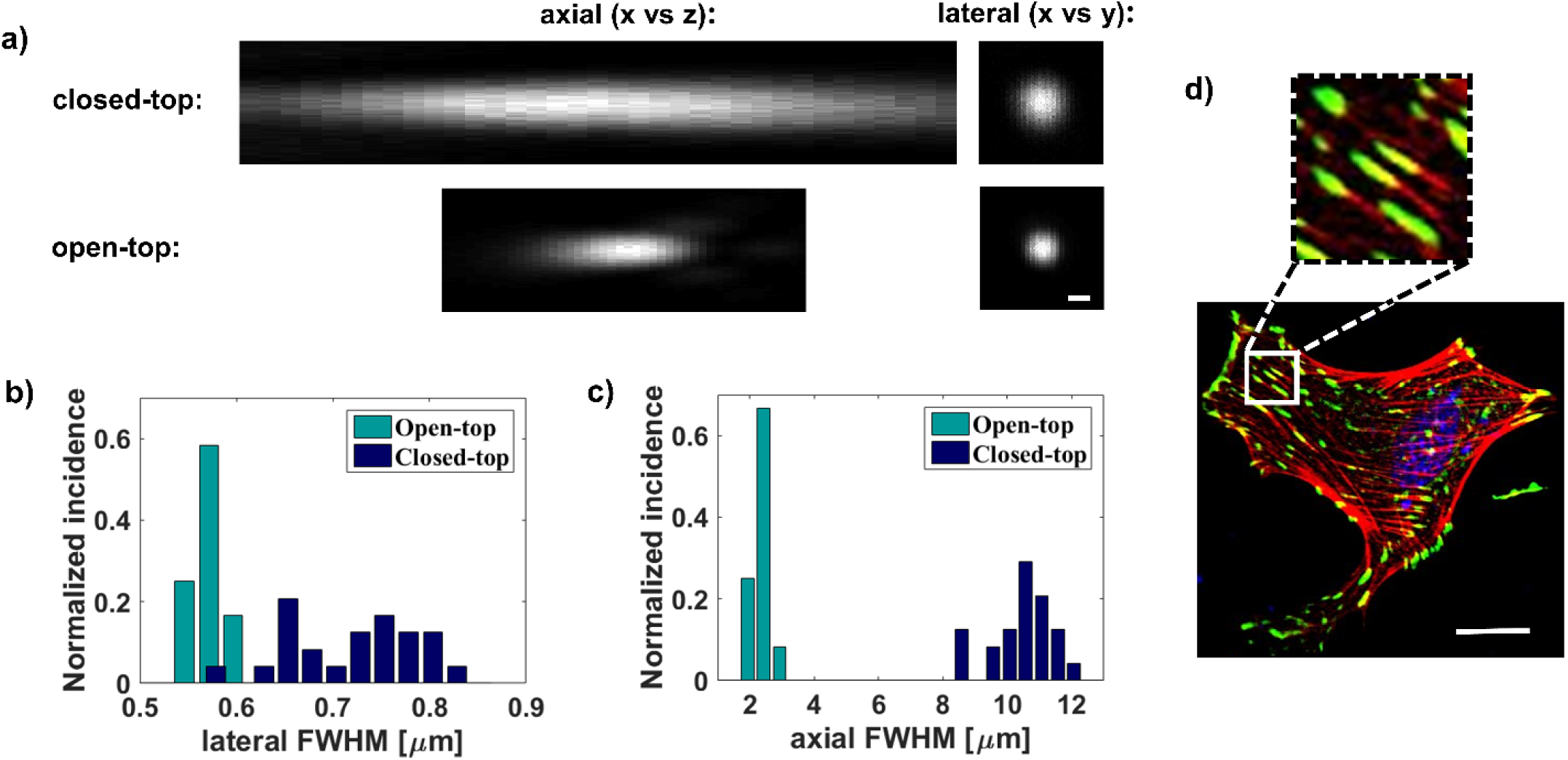
Imaging resolution improvement achieved by making the micro-stretcher open-top. **a)** Axial and lateral fluorescence profiles of a bead (diameter=200nm) imaged within the hybrid microstretcher made open-top and under a 800 μm thick PDMS layer. The scale bar is 500 nm. The distributions (n_beads_=24 for each condition) of lateral (**b**) and axial (**c**) FWHMs show significant loss of resolution (and increased variations) upon the introduction of the PDMS layer in the light path. The average lateral FWHM of the bead profile is 0.57 (s.d. = 0.02 μm) in the open-top case while it is 0.72 (s.d. = 0.07 μm) in the closed-top case, translating into 0.53 (s.d. = 0.02 μm) and 0.69 (s.d. = 0.07 μm) FWHM resolutions upon deconvolution of the bead size. d) Immunofluorescence composite image of a HFF cell adhered on the fibronectin-coated PDMS membrane of a hybrid microstretcher made open-top. The actin filaments are shown in red, the DNA in blue, and the vinculin in green. The inset magnifies a few focal adhesions and attached actin filaments. The scale bar is 25 μm. All the images were acquired with a laser scanning confocal microscope using a 25X objective.

### Cell mechanoresponses to repeated changes in strain direction

It was previously demonstrated that the reorientation response of cells subjected to two subsequent periods of cyclic stretching along perpendicular directions was qualitatively similar^31,45^. Prior to the present study, the question remained however whether the responses are quantitatively different and whether cellular realignment can be induced a larger number of times. Moreover, in both of these previous studies, only regions of approximately uniform strain (no strain gradient) were analyzed. Fig. 3a-f display normalized incidence histograms of the angle differences between the experimental cell normal orientations and the X-axis following seeding, and then 5 subsequent stretching periods (11 hours each), alternating between X- and Y-stretching. It can be observed that the cells are randomly oriented following seeding, and then they successfully realign perpendicularly to the stretching directions five successive times. Since the strain direction varies across the membrane, and to ease comparison from one reorientation to the other, Fig. 3g-l presents the same data as Fig. 3a-f but the orientations are reported with respect to the local principal strain direction. Moreover, considering that lower strain amplitudes are inefficient at inducing reorientation^10,26^, regions of sub-5% strain amplitude were excluded in Fig. 3g-l. As a result of these two considerations, a systematic narrowing is observed in the angle distributions from Fig. 3a-f to g-l. Interestingly, it can be seen from both sets of histograms that, although the cells do realign largely perpendicularly to the strain direction after each stretching periods, there is progressive broadening of the distributions (in Fig. 3, from a to f, and also from g to l). This reveals a decrease in cell normal alignment with the principal strain direction as time progresses, and thus as the stretching direction alternates.

**Figure 3.**
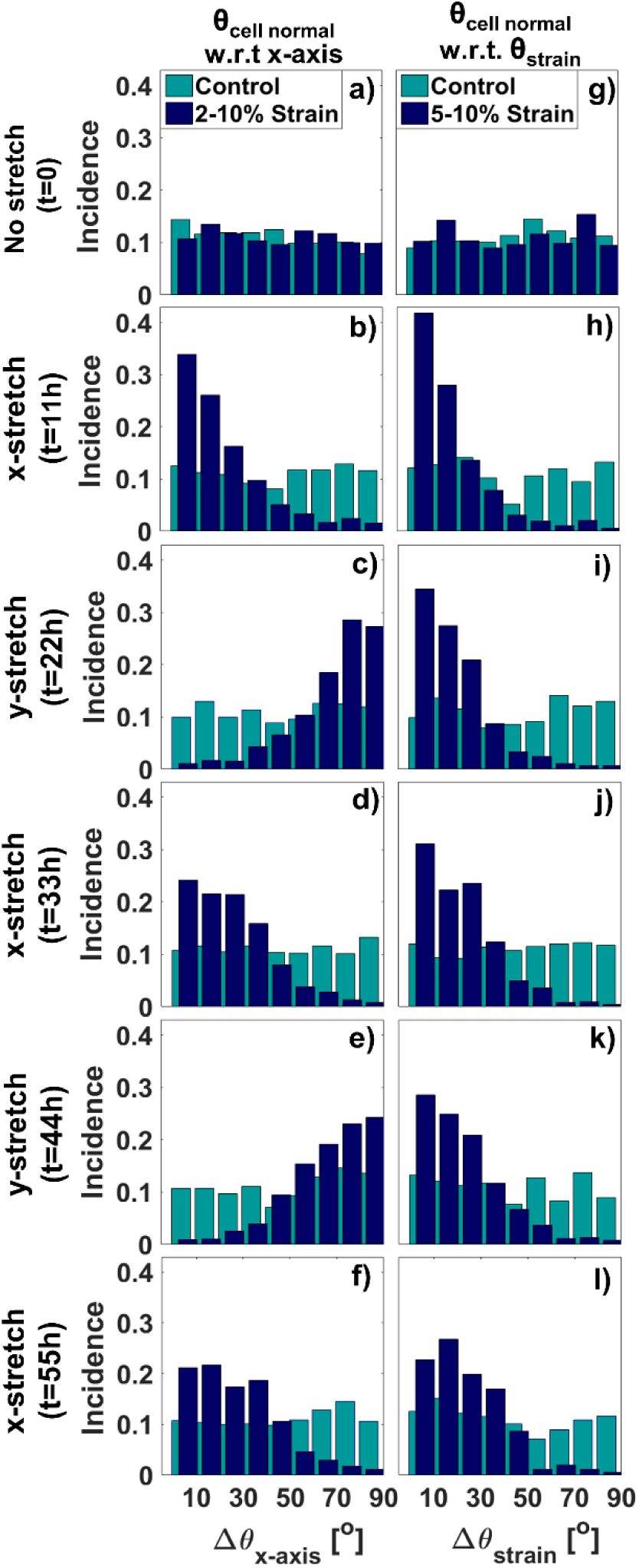
Reorientation analysis of HFF cells subjected to 5 changes in stretching direction over a period of 55 hours. **a-f)** Normalized incidence histograms of the angle differences between the experimental cell normals and the X-axis, considering the full membrane. **g-l)** Normalized incidence histograms of the angle differences between the experimental cell normals and the underlying maximum principal strain direction (considering the regions of high strain amplitude (> 5%)). The histograms **a)** and **g)** show the orientation distributions after 15 hours of seeding. The histograms **b-f)** and **h-g)** show the orientation distributions (of the same cells) after each of the five-successive cyclic-stretching periods of alternating directions (each lasting 11 hours). All the histograms comprise the combination of 6 unstretched control experiments as well as the combination of 6 cyclically stretched experiments. For **a-f)** n_control_ ≈ 1.4×10^3^ cells and n_strain_ ≈ 1.0×10^3^ cells while for **g-l)** n_control_ ≈ 8.5 ×10^2^ cells and n_strain_ ≈ 6.3×10^2^ cells.

The evolution of the cell orientations and morphologies was further analyzed and the results are reported in Fig. 4. The average angle differences between the X-axis and the experimental cell normals are displayed in Fig. 4a. The control is stable around 45° since the cells are randomly oriented and the data are reported between 0° and 90°. In contrast, the mean orientation of the cyclically-stretched cells with respect to the X-axis oscillates between low and high values. It is worth noting that the *means* are 20°-25° away from the *preferential* orientations (∼0° for X-stretching and ∼90° for Y-stretching, as seen in the histograms 3a-f) because of the statistical nature of the realignments^7^ and the fact that the data are reported between 0° and 90° (as opposed to between 0° and 180°). Fig. 4b shows the average angle difference between the local principal strain directions and the experimental cell normals, in regions of high strain amplitude (> 5%). The qualitative assessment that we noted previously is confirmed in this plot: the cell normal alignment with the principal strain direction decreases progressively. It is worth emphasizing that there is no residual oscillation in the data, meaning that complete strain-induced cell realignment was achieved after every uninterrupted stretching period. The near-linear increase of mean Δθ_strain_ in Fig. 4b, corresponding to a “poorer” alignment of the cell normals with the local strain directions, can be explained by the presence of another mechanical cue.

**Figure 4.**
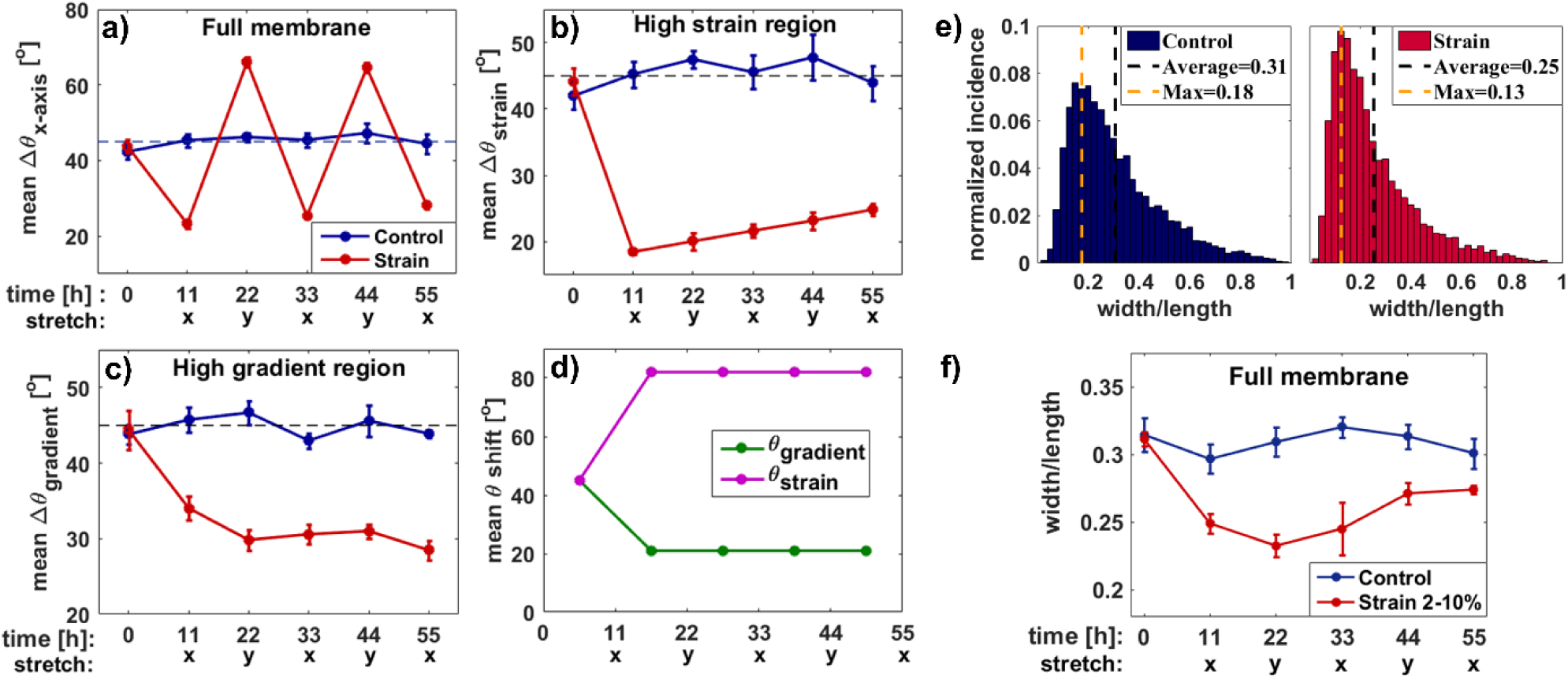
Evolution of cell orientation and elongation over the course of multiple reorientations. **a)** Mean angle differences between the experimental cell normals and the X-axis (mean Δθx-axis) as a function of time (and stretching direction), considering the full membrane area. ncontrol ≈ 1.4×10^3^ cells and nstrain ≈ 1.0×10^3^ cells. **b)** Mean angle differences between the experimental cell normals and the principal strain directions (mean Δθstrain) as a function of time (and stretching direction), considering the regions of large strain amplitude (> 5%). ncontrol ≈ 8.5 ×10^2^ cells and nstrain ≈ 6.3×10^2^ cells. **c)** Mean angle differences between the experimental cell normals and the principal strain gradient directions (mean Δθ_gradient_) as a function of time (and stretching direction), considering the regions of large strain gradient amplitude (> 7% mm^−1^). n_control_ ≈ 6.2 ×10^2^ cells and n_strain_ ≈ 4.6×10^2^ cells. In the plots **a-c)**, the blue curves show the unstretched control experiments and the red curves show the cyclically-stretched experiments, while the doted lines show the expected mean angle for perfectly randomly oriented cells. The errors on the means in **a-c)** are systematically lower in average for the stretched cells since the alignment is consistently guided by the strain field. **d)** Mean rotation angle that the cells would need to cover (mean Δθshift) - as a result from a change in stretching direction - if they were to perfectly align with the strain direction (green) or with the strain gradient direction (purple), in the regions of large strain amplitude and large strain gradient amplitude, respectively. Since the cells are randomly orientated before stretching, the first angle is 45°. **e)** Normalized incidence histograms of the width-over-length ratio for the unstretched control experiments (blue) and the cyclically-stretched experiments (red), with the data from the 5 successive reorientations combined together. n_control_ ≈ 6.8 ×10^3^ cells and nstrain ≈ 5.1×10^3^ cells. **f)** Mean width-over-length ratio of the cells as a function of time (and stretching direction), considering the full membrane. The blue curve shows the unstretched control experiments and the red curve shows the cyclically-stretched experiments. ncontrol ≈ 1.4×10^3^ cells and nstrain ≈ 1.0×10^3^ cells. In plots **a-c)** and **f)**, the error bars represent the standard error of the mean within the 6 unstretched control experiments and the 6 cyclically-stretched experiments. The data points associated with 11 h and 55 h are statistically different in **a), b), c)**, and **f)**, with p-values of p=0.011, p=0.0004, p=0.016, and p=0.016, respectively.

The strain gradient has recently been identified as a genuine mechanical cue involved in the reorientation response of fibroblasts^26^ and it can indeed explain the trend observed in Fig. 4b. The mean angle difference between the experimental cell normals and the principal strain gradient directions is displayed in Fig. 4c, considering the regions of larger strain gradient amplitude (> 7% mm^−1^). Interestingly, conversely to the cell normal alignment with respect to the strain, there is a slight improvement of the cell normal alignment with respect to the local strain gradient direction, as the experiment progresses. Since the improvement of one is on the same order of magnitude as the deterioration of the other (and the two membrane regions considered overlap greatly), we speculate that these two trends result largely from an interplay between the cellular mechanoresponse to these two cues (strain and strain gradient fields).

To understand how a progressive improvement in cell normal alignment with the gradient is possible despite the drastic changes in the applied stretching direction (X or Y) imposed every 11 hours, the average local change in membrane’s gradient direction is evaluated (Fig. 4d). In average, in the regions of large gradient amplitude considered in Fig. 4c, the direction of the strain gradient shifts by only ∼20° for every change in stretching direction (due to the specific geometry of the strain field). Since this angle is relatively low, the trend observed in Fig. 4c could be explained by the fact that cells take longer to align with the strain gradient that they do with the strain itself. Their alignment progression over each stretching period is not abolished because the gradient direction only changes by 20° every 11 hours. In other words, they can slowly adapt even though the gradient response seems slower. In contrast, we determined from our extracted strain field that the direction of the strain shifts in average by ∼ 80° between every change in stretching direction (in the regions of large strain amplitude considered in Fig. 4b). This angle is relatively high and as we will explicitly demonstrate in Fig. 5, 11 hours is sufficient for the cells to complete the strain-induced reorientation response.

**Figure 5.**
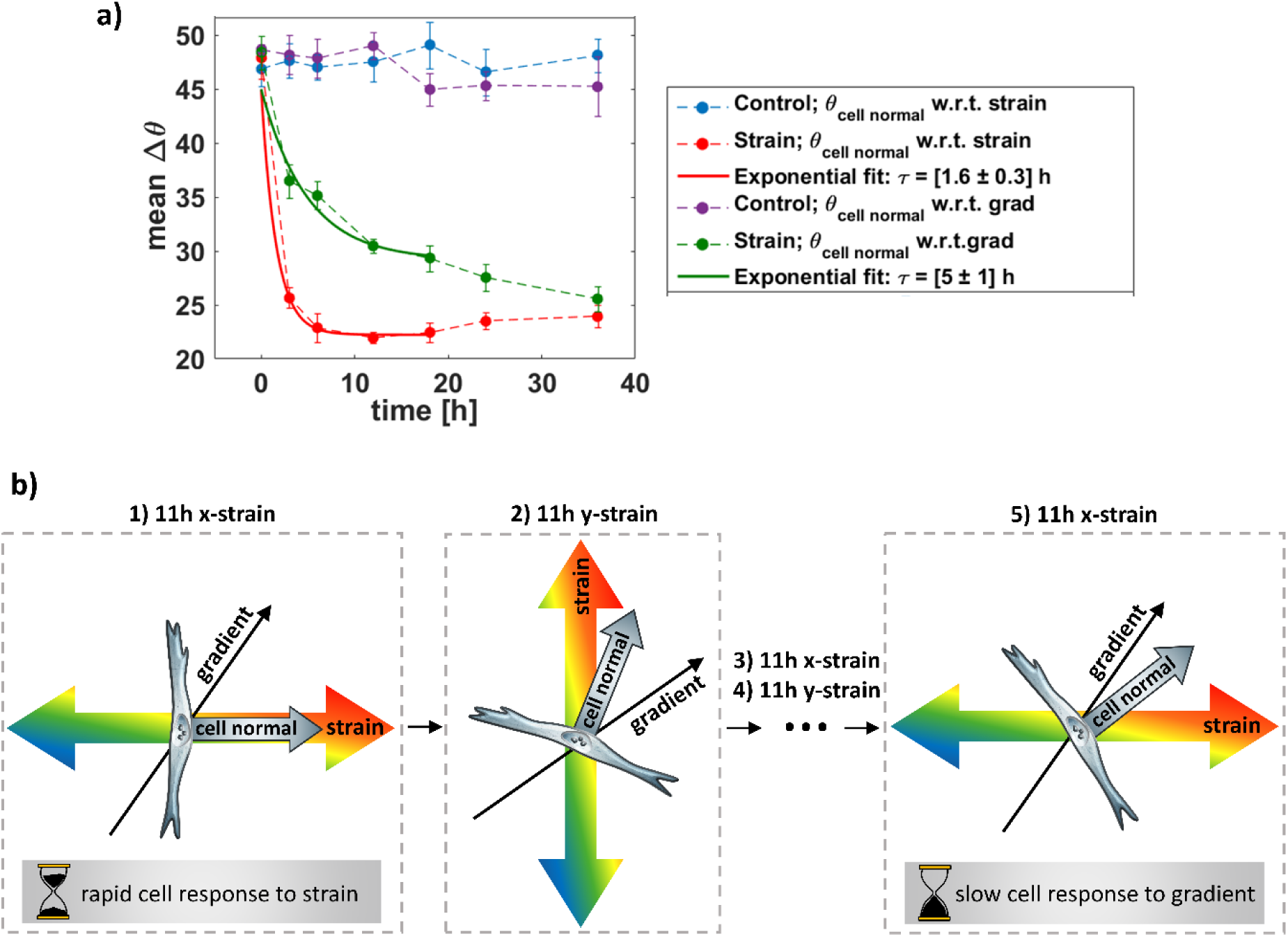
Time-scale mismatch between the strain and strain gradient responses. **a)** Time-dependence of the cellular response over 36 h of X-stretching. Temporal evolution of the mean angle differences between the experimental cell normals and either the strain directions (red curve) or the strain gradient directions (green curve) in the regions of large strain gradient amplitude (> 7% mm^−1^). Exponential fits are also displayed, considering only the first 18 hours to ensure near-constant cell density (<20% change). The error bars on the experimental points represent the standard error of the mean within the 5 unstretched control experiments and the 8 cyclically-stretched experiments. The uncertainty on the exponential fits represent the standard error of the mean of the time constants extracted for the individual experiments. n_control_ ≈ 6.5 ×10^3^ cells and nstrain ≈ 8.4×10^3^ cells. The rise of the red curve beyond 12 h is statistically significant (p = 0.046, data of the last two time points compared to the data of the two previous points). The drop of the green curve beyond 12 h is statistically significant (p = 0.011, data of the last two time points compared to the data of the two previous points). **b)** Evolution of the cellular orientation in response to complex and varying strain field over 55 hours. 1) After 11 hours of cyclic X-stretching, the cell normal is well aligned with the principal strain direction (not with the gradient direction). 2) 11 hours after changing the stretching direction, the cell normal is mostly aligned with the strain direction, but the effect of the gradient is starting to influence the cell direction. 3-4) Over the subsequent changes of stretching direction, the cell normal continues to further align with the gradient direction and less with the strain direction. 5) 11 hours after the fifth change in stretching direction, the cell normal is approximately halfway between the gradient and the strain directions. This pattern of cellular reorientation can be explained by two elements. First, the geometry of the complex strain field is such that in average, each shift of stretching direction consist in a ∼80° change of principal strain and a ∼20° change of gradient direction. Second, the cell reorients more rapidly in response to strain than to the strain gradient.

The effect of cyclic stretching on the cell morphology is examined in Fig. 4e where the distribution of aspect ratios is presented for non-stretched and stretched cells. It can be seen that cyclic stretching causes cell elongation, in agreement with previously reported studies^7,8^. Interestingly, we observe a narrowing of the distribution for the stretched cells. Fig. 4f displays the evolution of the cell morphology over the course of the multiple reorientation experiments. Except for the first-time point (at t=0 hour), for which the control and strain data overlap, the stretched cells are systematically more elongated that the control ones. Moreover, the stretched cells’ morphology appears to further evolve following the initial elongation. We observe a significant recovery of the aspect ratio from 11 to 55 hours (p = 0.016, see caption Fig. 4f), once again suggesting a rather complex mechanoresponse that involves possibly more than one time-scale.

### Time-dependent cellular responses to mechanical cues

The characteristic cell behaviors observed in the multiple reorientation experiments (Fig. 3 and 4) suggest the presence of a dual time-scale mechanoresponse which we now seek to explicitly investigate. Fig. 5a presents the temporal evolution of the cell orientations over 36 hours of X-stretching (no change in stretching direction) in the region of high strain gradient amplitude (> 7% mm^−1^). The cell alignment with the strain direction (red) and the strain gradient direction (green) are both presented by showing the mean angle difference (mean Δθ) between the cell normal and the strain or the strain gradient directions, respectively. To quantitatively assess the temporal evolution of the cellular response to the strain field, exponential decay fits (Δθ = A[e^−t/τ^ - 1]+45°) were performed resulting in time constant of τstrain = [1.6 ^±^ 0.3] hours and τ_gradient_ = [5 ^±^ 1] hours. Note that the last two time points were excluded from the fits to ensure that the cell density was approximately constant (<20% change).

The mean Δθ_strain_ initially drops abruptly, indicating a rapid realignment response with the strain direction, which is in qualitative agreement with previous studies^15,46^. It then reaches a minimum between 10 and 20 hours, after which it undergoes a slight increase, indicating a reduced alignment (p = 0.046, see caption Fig. 5). As suggested above, this rise can be explained by the concurrent increase in cell alignment with the strain gradient direction (p = 0.011, see caption Fig. 5). Since the mean Δθ_gradient_ exhibits a much slower drop, the relative importance of the gradient cues with respect to that of the strain cues only becomes significant at later time points. Consequently, as the cell normal alignment with the gradient direction continues to improve after 20 hours, its alignment with the strain direction is worsen.

Taken together, our results demonstrate a consistent cellular response to given mechanical cues (strain and strain gradient) despite drastic changes in strain direction, suggesting the absence of apparent major stretched-induced irreversible changes. They also reveal that the mechanoresponses associated with two different aspects of an applied strain field take place on different time scales. This suggests that there might be subtle differences in mechanotransduction mechanisms involved in the cellular responses to strains and strain gradients. It has been suggested in our previous study that the strain gradient sensing mechanism possibly involves the creation of an intracellular gradient of biological signals^26^. This could explain why the mechanoresponse of cells to the strain gradient takes place on a longer time scale. In this study, the cellular responses depend on the specific history of the mechanical cues: in the presence of non-uniform and time-varying strain fields, the interplay between cues can give rise to a complex cellular mechanoresponse with a temporal evolution that depends on previous events.

## CONCLUSION

We presented a microfluidic stretcher array developed to study the reorientation response of human fibroblasts in a non-uniform strain field subjected to repeated changes in stretching directions. Its improved capabilities and usability with respect to previous micro-stretcher designs make it a device of choice for future cyclic- or static-stretching studies aiming to investigate the effect of non-uniform strain fields. Using this device, we exposed the adaptability of cells to discontinuous mechanical cues by achieving multiple successive cellular reorientations. By demonstrating that two aspects of the strain field drive cellular responses on different time-scales, we gained further insight on the physical interaction between a cell and its microenvironment. Such strategy of exposing cells to increasingly complex in vitro environment is key to understanding the interplay of different cues and how it affects the cells’ fate.

## MATERIALS AND METHODS

### Microfluidic stretcher design and fabrication

#### SU-8 master fabrication

Fig. 6 describes the device and its general fabrication strategy. PDMS microdevices were produced using a replica molding approach. To fabricate the master mold, a photomask pattern (Fig. 6a) was transferred to negative SU-8 photoresist (Microchem, MA, USA) using standard soft lithography procedures on a silicon wafer. The thickness of the pattern (340 μm) involved the deposition of two separate layers of 2050 SU-8 (2×170 μm). Standard cleaning, prebaking, spin coating, soft baking, UV-exposure through the photomask, post-baking, developing, and hard baking procedures were employed (consistent with manufacturer’s recommendations). Note that the design includes deep (340 μm) and narrow (120 μm) trenches between the cell chambers and the vacuum chambers (Fig. S1 of the SI), making conventional SU-8 development procedures insufficient. Instead, we used a more efficient strategy which is presented in the SI.

**Figure 6.**
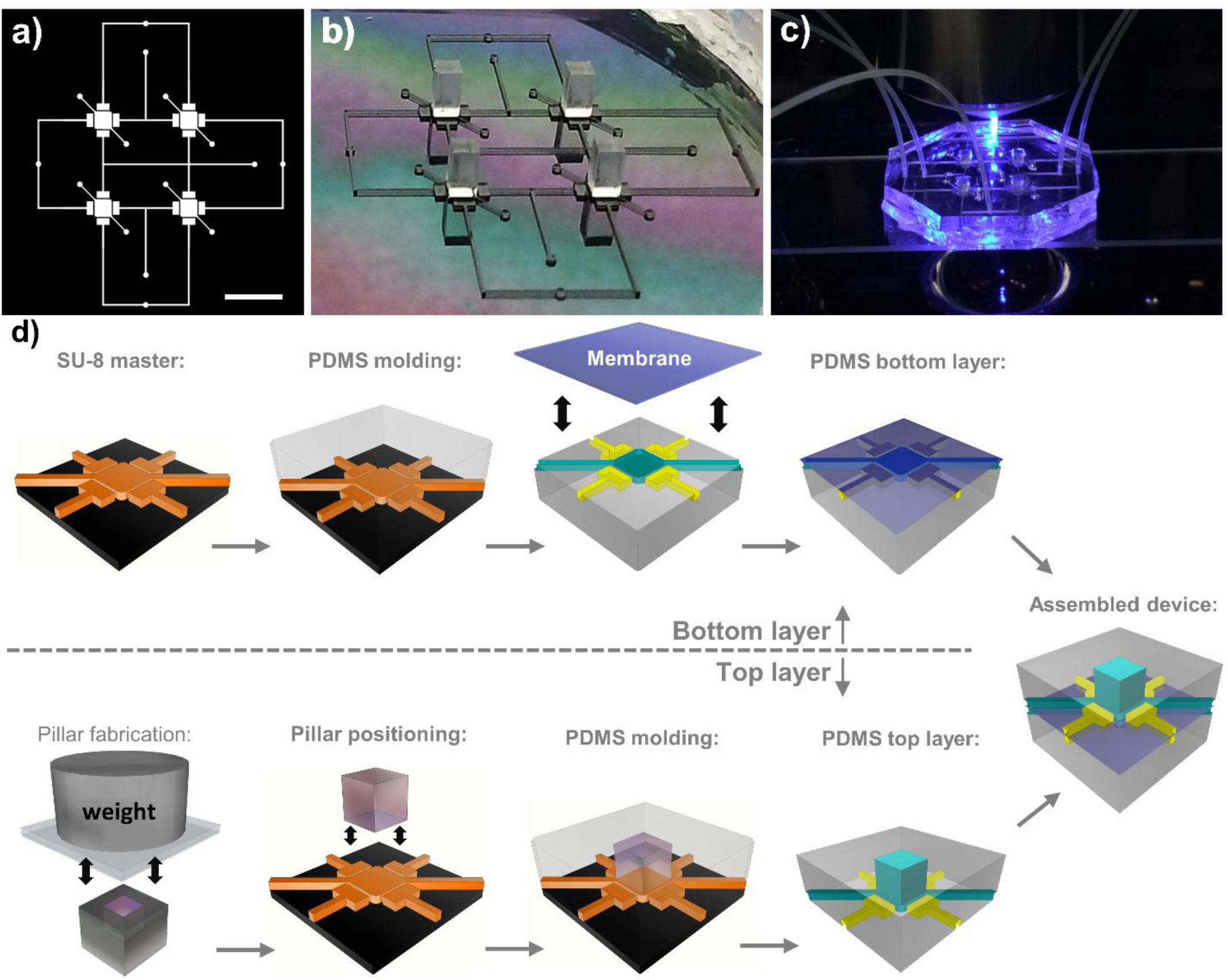
Microfluidic device and general fabrication strategy. **a)** Photomask of the top PDMS layer, showing the microdevice’s pattern which includes the microfluidic loading channels, the vacuum pumps, and the cell chambers. The scale bar is 5 mm. **b)** Picture of the photoresist master used to mold the top layer of the device, after positioning of the PDMS pillars. **c)** Picture of the hybrid microfluidic stretcher during high resolution fluorescence imaging. **d)** Schematics of the main fabrication steps (showing one of the four cell chambers), which involve in particular the fabrication of chemically treated PDMS pillars (See SI3). The microfluidic device comprises three PDMS layers: two micropatterned layers and one thin (10 m) membrane, which is assembled on the PDMS bottom layer.

#### Augmented master combining SU-8 and PDMS for molding the top layer

To expand the design possibilities, we combine this master with movable chemically-treated PDMS sub-molds (pillars in our case, see SI for fabrication method). Specifically, the movable PDMS sub-molds are straightforwardly positioned onto the desired master features (on the cell chamber in our case), under a microscope (see Fig. 6b). The chemical treatment of the PDMS sub-molds prevents their polymerization with the uncured PDMS to be molded. Interestingly, it also facilitates the precise positioning of the sub-mold onto the SU-8 feature by providing an excellent balance between adhesion (to ensure that it stays in place when pouring uncured PDMS) and slipperiness (to ease fine repositioning). In practice, it is easy to achieve sub-100 μm positioning resolution with tweezers only, but higher precision tools (mask aligner or micro-positioners) can be used if greater resolution is needed. The details of the PDMS pillars fabrication method and chemical treatments are provided in the SI. We note that an alternative method to achieve PDMS features of different heights would be to fabricate a multi-layers master via strategic photoresist layering and UV exposition patterning steps^47,48^. However, the SU-8 feature heights achievable (and thus PDMS open-top features) are typically limited to a few hundred microns and it becomes increasingly challenging with the number of different heights. Producing mm-thick PDMS layers is often desirable in order to ensure structural integrity upon manipulations and connection to external tubes. In contrast, the PDMS sub-mold strategy we used and outlined above has no such limitations (maximum height and number of different heights) in addition to be fast and simple, making it a method of choice for complex cell-scale device features. A summary of the other methods that we considered for the PDMS top layer fabrication is presented in the SI.

#### Device molding and assembly

The PDMS (Sylgard 184 silicone elastomer) top and bottom layers were molded using the microfabricated master described above. They were aligned and bounded using an air-plasma system (Glow Research, Tempe, AZ, USA) and a mask aligner (OAI, CA, USA). Prior to assembly, the bottom layer was bonded to a 10 μm thick PDMS membrane, which was spin coated onto a clean silicon wafer. The top layer is composed of features having two significantly different heights (∼2 mm for the cell chambers, and 340 μm for other features). Uncured PDMS was then poured onto the augmented master mold described above (SU-8 features and positioned pillars) until PDMS barely covered the pillars.

### Cell culture, stretching experimental procedures, and imaging

The cells were cultured in a standard incubator (37°C, 5% CO_2_) and kept in Dulbecco’s Modified Eagle’s Medium (DMEM) with 10% FBS and 1% streptomycin/penicillin added (Hyclone Laboratories, Logan, UT, USA). Each microdevice was air-plasma treated (Glow Research, Tempe, AZ, USA) for 5 minutes at 70 W for hydrophilicity and to reduce the risk of trapping air bubbles. Flushing the microchannels with 70% ethanol for 5 minutes and then with autoclaved phosphate buffered saline (PBS) solution ensured sterilization and removal of ethanol from the microdevice, respectively. Prior to cell seeding, in order to promote the cell adherence to PDMS, each membrane was functionalized via incubation for 4 hours in a solution of fibronectin 10 μg/ml of HEPES-buffered salt solution (HBSS; 20 mM HEPES at pH 7.4, 120 mM NaCl, 5.3 mM KCl, 0.8 mM MgSO4, 1.8 mM CaCl2 and 11.1 mM glucose). A density of adhered cells of ∼300 cells/membrane was achieved by injecting a solution of culture medium with 5×10^4^ cells/ml and using an incubation time of 15 hours. Note that two different seeding methods can be employed with our device. First, the cells can be loaded into the device’s stretching chamber via the microfluidic channels using pressurized vials. Second, a micropipette can be used to directly deliver the cell solution if the PDMS layer acting like a top sealant has been peeled off beforehand (which is the case when the microfluidic properties are not required during the experiment, see section Result and Discussion).

All the stretching experiments were carried out within the incubator to ensure optimal environmental conditions. The vacuum pumps were controlled via a Labview program creating sinusoidal stretching waveforms (2 to 10% strain amplitude across the membrane, 0.5 Hz). To acquire the cell orientation data over the course of the experiments, the cells were imaged before and between stretching periods by interrupting the cyclic stretching momentary (∼10 minutes). A Nikon TiE inverted phase contrast EPI microscope (long working distance 10X objective) was employed to image the living cells within the devices. Immediately after imaging, cyclic stretching was reinitiated to continue the experiments. Fig. 2d was obtained in separate experiments by immunofluorescence staining and imaging the cells directly within the microfluidic device, 15 hours after seeding. Details of the staining protocols are given in the SI.

### Membrane strain field calculation

The detailed method to extract the strain field was published previously^26^. First, fluorescent beads (FluoSpheres, 200 nm, Invitrogen, CA, USA) embedded in the elastic membrane were repeatedly imaged with an EPI fluorescence microscope (Nikon, Tokyo, Japan) as the membrane was stretched. Their displacements were then extracted with a custom Matlab script in order to calculate the maps of the Green strain matrix elements, and subsequently the maps of the maximum (**ε1**(x,y)) and minimum (**ε2**(x,y)) principal strain fields^49^. The interpolated and boxcar smoothed strain fields of 5 devices (and their mirror images, since the device is symmetrical) were averaged together to provide an accurate representation of the device population. For every position across the membrane, the maximum principal strain field is associated with the direction of maximum stretch while the minimum principal strain field is associated with the direction of minimum stretch or maximal compression. In our device, the amplitude of the former largely dominates that of the latter across the full membrane. Therefore, in the orientation analysis, the cell normals were reported with respect to the maximum principal strain direction when specified as Δθstrain. Finally, the gradient field (of the maximum principal strain was calculated everywhere across the membrane.

### Cell orientation and elongation analysis

The background noise was reduced from the phase contrast images with a Fast Fourier Transform bandpass filter using ImageJ. To capture the cell morphologies, binary images were generated with the adaptive threshold plugin. The cell orientations, positions and elongations were obtained via the Analyze Particles tools of ImageJ. The analysis of the extracted cell data was achieved with a custom Matlab script. In particular, the orientation of each cell normal was reported with respect to either the X-axis (Δθ_x-axis_), the local strain direction (Δθ_strain_), or the local strain gradient direction (Δθ_gradient_). The cells located within 150 μm of the chamber’s walls were not considered to eliminate possible edge effects. At the end of the analysis, all angles were reported between 0 and 90° in order to produce Fig. 3 to 5.

### Statistical analysis

Statistical significances were determined using unpaired two-tailed t-test, assuming unequal variances. The number of samples and the number of independent experiments are specified in each figure. The t-test were performed by considering the averages extracted from the independent experiments (as opposed to considering every single cell orientations from all experiments). The error on the degradation of the average lateral FWHM represents the standard error on the mean.

## ACKNOWLEDGEMENTS

This work was supported by individual Natural Sciences and Engineering Research Council (NSERC) Discovery Grants to M.G. and A.E.P. S.C-L. was supported by NSERC Postgraduate Scholarships-Doctoral (PGS D). M.G. acknowledges support from the Ontario Ministry of Research and Innovation, and the Canada Foundation for Innovation (CFI). A.E.P. gratefully acknowledges generous support from the Canada Research Chairs (CRC) program.

