## Supplementary Materials for "Time dependence of cellular responses to dynamic and complex strain fields"

**SI1: Additional details about the microfluidics device**

The microfluidics device was designed in such a way as to enable the simultaneous generation of biologically relevant strain and strain gradient amplitudes (varying from 2% to 10%, and from 0%$mm^{-1}$ to 14%$mm^{-1}$, respectively). For a given non-uniform strain pattern, the achieved gradient amplitudes scale inversely with the device dimensions. Therefore, it would be possible to generate higher gradient amplitudes without increasing the strain amplitude (~10% being known as a cell responsive amplitude range[1–3]), simply by further scaling down the microdevice pattern. To study different cell mechanisms under strain gradients, microstrecher devices are thus better suited than their macro counterparts. Using arrays of cell chambers allows for high throughput fabrication and experiments, and their currently partially coupled channels could easily be re-designed for independent control if needed.

**SI2: Photoresist development procedure for the master mold fabrication**

Clean, uniform, and fully developed photoresist trenches could not be obtained by using manual shaking (or ultra-sonic bath) of the solution during the development process. We used a simple, efficient, and low-cost strategy to overcome difficulties associated with the thickness of the SU-8 layers (see Fig. S1b). We simply attached the master mold wafer to a stirring bar and immersed it in a beaker filled with developer solution. The speed of the stirring plate was set to 400 rmp. This method was found to be gentle on fine SU-8 structures (and thus ensure features' integrity) while creating a fast-rotational motion of the solution such as to maximize the development process.

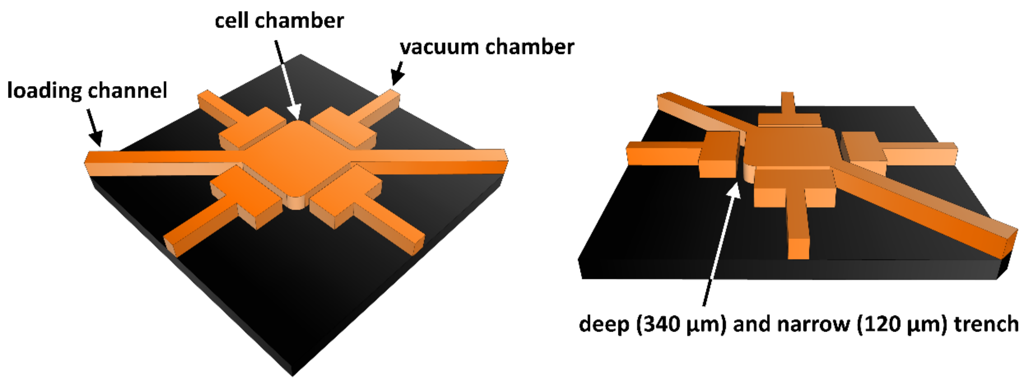


**Figure S1. Microfluidics SU-8 master mold.** Representation of the SU-8 master mold (cropped view) showing the different parts of the design. The deep trenches between the cell chamber and the vacuum chambers require a special development procedure.

**SI3: PDMS pillars fabrication method**

In order to fabricate the PDMS pillars, micro-milled aluminum molds were employed. Two important designing aspects need to be considered. First, the PDMS shrinks[4] during the curing process (with our conditions, we increased all three pillar dimensions by ~1.5% to compensate for shrinking). Second, it is advantageous to include error margins to loosen the centering/positioning requirements onto the SU-8 features of the master (we decreased the pillar depth and width by ~10%). To fabricate the PDMS pillars from the aluminum mold, uncured PDMS was simply poured into them and then degassed in a vacuum chamber. To ensure the production of pillars having all six surfaces flat, the top surface of the mold was then closed. A non-frosted acetate sheet was positioned directly onto the uncured PDMS (to facilitate demolding) after which a glass slide and a 500 g weight were placed on top of the acetate sheet to ensure a flat surface and to expelled extra PDMS, respectively (see Fig. 6c of the manuscript). After curing and demolding, the PDMS pillars were chemically treated to prevent them from bounding to uncured PDMS during the fabrication of the device's top layer. This chemical treatment consists in i) an air-plasma treatment (70W, 5 min), ii) two hours of soaking in methanol with 2% trichloro(1H,1H,2H,2H-perfluorooctyl)silane (Sigma-Aldrich) and iii) one hour of heating at 100°C to ensure solvent evaporation.

**SI4: Additional discussion on the advantages of using an open-top device for fluorescence imaging.** In the most common cases of oil-immersed objectives or water-dipping objectives, near-diffraction limited resolution requires the refractive index (n) to be approximately constant between the objective and the sample, namely ~1.52 or ~1.33, respectively. In both cases, the introduction of a PDMS layer (n≈1.42)[5] is thus highly detrimental. Certain water-immersion objectives have correction rings which can compensate for the introduction of glass coverslips up to 0.17 mm thick and which could partially compensate for the spherical aberrations introduced by a PDMS layer. However, the latter are generally too thick and more importantly too non-uniform in practice. Non-uniformities in the thickness or density of even a thin refractive index mismatched layer result in image deterioration[6,7]. Moreover, water-immersion objectives with correction rings are significantly more expensive than the other types mentioned above. For this reason, and because complex staining procedures strongly benefit from having an open access to the cell chamber, the alternative of using a permanently closed glass-top microstretcher, rather than an all-PDMS device with removable top, is not optimal in many contexts.

**SI5: Additional details about the staining and imaging procedures**

The microfluidics top membrane (which acts like a lid to close the cell chamber) was peeled off prior to performing the staining and imaging procedures. The cells were fixed using 3.5% paraformaldehyde for 10 minutes and permeabilized with 0.5% TritonX-100 for 3 minutes. Phalloidin conjugated to Alexa Fluor 546 (Invitrogen) and DAPI (Invitrogen) were respectively employed to stain the actin filaments and the DNA. Vinculin was stained with a monoclonal mouse anti-vinculin antibody and a rabbit anti-mouse IgG secondary antibody conjugated to Alexa Fluor 488 (Invitrogen). Details of the staining procedures have been previously published[8,9]. Minor modifications were required to take into account the fact that cell staining is performed within the microdevices. The incubation time of every staining and washing (PBS solution with 5% horse serum) steps was doubled. This procedure was found to compensate for the low fluid mixing taking place in the cell chamber, as well as for the loss of photons arising from PDMS scattering and surface reflection during fluorescence imagining. After staining, the cells were immersed in cold PBS solution and imaged with an upright laser scanning multiphoton confocal microscope (Nikon A1RsiMP) with a long working distance 25x objective (NA=1.1).

**SI6: Alternative methods to fabricate the top PDMS layer**

**Table S1.** Fabrication methods investigated to produce varying feature heights within a single PDMS layer.

| **Fabrication methods investigated** | **Pros** | **Cons and fabrication difficulties** |
|---|---|---|
| 3D printed mold to replace the SU-8/Si master mold | • simple<br>• no height limitations | • typically lower resolution<br>• material not necessarily compatible for PDMS molding |
| Strategic photoresist layering and UV exposition patterning | • retains high resolution | • heights achievable typically limited to a few hundred microns<br>• increasingly challenging when the number of different heights needed augments |
| Fabricating two distinct SU-8 masters (in order to fabricate two distinct layers instead of one with multiple feature heights) | • retain high resolution | • heights achievable typically limited to a few hundred microns<br>• requires additional alignment and plasma bounding steps involving thin (easily-deformable and damageable) layers; |
| Post-fabrication punch-holes | • Simple and fast | • low resolution<br>• generally limited to circular holes<br>• compromises the layer's integrity and leaves residues on the hole's edge<br>• impossible to leave a peelable membrane |
| Post-fabrication laser ablation | • high flexibility with respect to the shape and depth ablated<br>• high resolution possible | • requires a complex positioning and monitoring setup as well as access to an appropriate laser [PDMS is transparent over a large range of wavelengths[10]; it would require either a UV laser (<300 nm) for single-photon absorption, or a longer wavelength femtosecond laser (near-infrared or visible) for multiphoton absorption]<br>• potential melting damages compromising the integrity and imaging capabilities |
| Sub-molds (PDMS, metal, or other) glued to the SU-8 features | • no height limitations | • high possibility of damaging the mold with the introduction of glue<br>• all glue tested had poor adhesion with the SU-8<br>• potential toxicity (glue residues) for the cells |
| Chemically-treated PDMS sub-molds positioned on the SU-8 features | • simple, cheap, and fast<br>• high success-rate<br>• no height limitations; | • more challenging when higher resolution (~sub-50 um) features are desired |
